# The mechanoreceptor PIEZO1 is a novel oncogene in glioma by promoting astrocyte reactivity

**DOI:** 10.1101/2024.04.30.591858

**Authors:** Pablo Blanco-Carlon, Miguel Ángel Navarro Aguadero, Pedro Aguilar-Garrido, Álvaro Otero-Sobrino, María Hernández-Sánchez, Andrés Arroyo-Barea, Osvaldo Graña-Castro, Ricardo Gargini, Rosa María García-Martín, Aurelio Hernández-Laín, Manuel Pérez-Martínez, Jesús Gómez Alonso, Sagrario Ortega, Joaquín Martínez-López, Miguel Gallardo, María Velasco-Estévez

## Abstract

Glioblastoma is the most common and aggressive brain tumour in adults. Despite advances in its molecular characterization, there is a gap-of-knowledge in the identification of *bona fide* drivers and potential therapeutic targets that could change the clinical picture. Mechanoreception, the sensing of mechanical cues by cells, has proven to be an important factor in cell biology, particularly in cancer. Piezo1 is a mechanoreceptor found in all cells and seems to play a role in different cancer types, such as gastric, breast or lung cancer. However, there is still lack of understanding about its role in the onset and progression of glioblastoma. Here, we show that Piezo1 acts as an oncogene in glioma by potentially promoting chronic astrocyte reactivity. We developed a novel transgenic murine model of Piezo1 overexpression in astrocytes. These animals had a significant reduction in overall survival and developed glioma with a penetrance of 30%. We also developed a PIEZO1-overexpressing U251 cell line and found that it had a more aggressive and reactive-like phenotype. Finally, we correlated the levels of PIEZO1 with the clinical outcome of a cohort of glioblastoma patients and observed that PIEZO1 is a biomarker of worse prognosis. However, PIEZO1 only correlated with worse prognosis in male patients, suggesting a sexual dimorphism. In conclusion, we identified Piezo1 as a *bona fide* driver of glioma, revealing its implication in astrocyte reactivity and identifying it as a biomarker for glioma in the clinic.

## Introduction

Glioblastoma multiforme (GBM) is one of the most common types of malignant brain tumor in adults, with an incidence of 2-3 per 100.000 adults worldwide^1^. Currently, the 2021 WHO classification of central nervous system tumors defines GBM as an isocitrate dehydrogenase (IDH)-wildtype grade IV astrocytic tumor, with IDH-mutant astrocytoma being considered an independent type^2^.

GBM patients have a median overall survival of approximately 15 months and an overall 5-year relative survival rate of 4–5%^3^. Despite advances in the molecular and behavioural characterization of GBM, there have been only moderate improvements in survival rates for patients with this disease in recent decades^4^, and new therapeutic options have limited ability to alter the course of disease. Therefore, the current standard of care for GBM patients, which is based on surgery followed by field radiotherapy combined with temozolomide chemotherapy^5^, has hardly changed over the last 30 years. All these features, together with its high recurrence rate (75-90%)^6, 7^ make it one of the most aggressive cancers.

Some of the main obstacles to achieving an effective therapeutic approach for GBM are its molecular heterogeneity and the absence of an identified *bona fide* tumor driver, which could be used as a therapeutic target. These features have also hampered the development of efficient GBM mouse models that mimic the microenvironmental and genomic characteristics of human brain tumours, with most current models involving more than two gene mutations or presenting a low GBM incidence^8^, which presents an added difficulty for research on this tumor.

Despite the heterogeneity of GBM, there are some hallmarks common to all tumours, such as neuroinflammation. GBM tumours are characterized by high levels of infiltrating immune cells and inflammatory cytokines, such as IL-6, IL-8 or CSF1, and there is strong evidence that the greater the inflammatory microenvironment is, the worse the prognosis^9–11^. Moreover, the activation of some inflammatory pathways in GBM cells, such as the STAT3 pathway, has also been associated with tumor progression^12, 13^.

Piezo1, also known as FAM38A, is a mechanosensitive nonselective Ca^2+^ channel that belongs to the PIEZO family. It was first identified and characterized in 2010 in a mouse neuroblastoma cell line by Coste and colleagues^14^. This channel is composed of more than 2500 acids, with a large number of transmembrane regions, and its structure consists of a homotrimer that forms a three-blade propeller plus an extracellular cap^15^.

Since its discovery, Piezo1 has been reported to play a role in a wide variety of cancers, including GBM. According to Chen et al., Piezo1 overexpression is associated with poor prognosis in GBM patients, and it increases the proliferation and aggressiveness of *D. melanogaster* gliomas^16^. Similarly, Piezo1 expression is positively correlated with ECM organization, cell adhesion, angiogenesis, cell migration and proliferation in human GBM tumours, making it a potential predictive marker in this disease^17^. Although the molecular mechanisms by which Piezo1 is involved in GBM biology remain unknown, it has been reported that this mechanoreceptor is important in the biology of healthy astrocytes, playing a key role in promoting astrocyte reactivity and cytokine release, thereby exacerbating neuroinflammation^18^, which could explain the association between Piezo1 and GBM. Nevertheless, there are many unresolved questions about the exact role that Piezo1 plays in GBM and the mechanism by which the mechanoreceptor affects GBM biology, so further investigation is needed to evaluate the potential of Piezo1 as a therapeutic target in GBM.

In this study, we elucidate the role of Piezo1 in the onset and progression of glioma. We developed a novel mouse model in which Piezo1 is overexpressed in astrocytes and found that this overexpression produces neuroinflammation and glioma. Furthermore, we studied the molecular mechanisms involved in the aggressiveness of GBM by developing and analysing the U251 cell line overexpressing PIEZO1. In conclusion, for the first time, we describe Piezo1 as a driver oncogene for glioma.

## Results

### PIEZO1 is a marker of poor prognosis in glioblastoma patients

To determine the impact of PIEZO1 expression in human GBM, we analysed 63 brain biopsy samples from GBM patients. Samples were classified as having low PIEZO1 or high PIEZO1 expression (*Figure 1A*). We observed that patients with high PIEZO1 expression presented a significant reduction in overall survival compared to patients low PIEZO1 expression (448 days in the high PIEZO1 group vs 616 days in the low PIEZO1 group, *p=0.03). Notably, there were no significant differences in the mean age at diagnosis between the two groups (*Figure 1C*).

**Figure 1:**
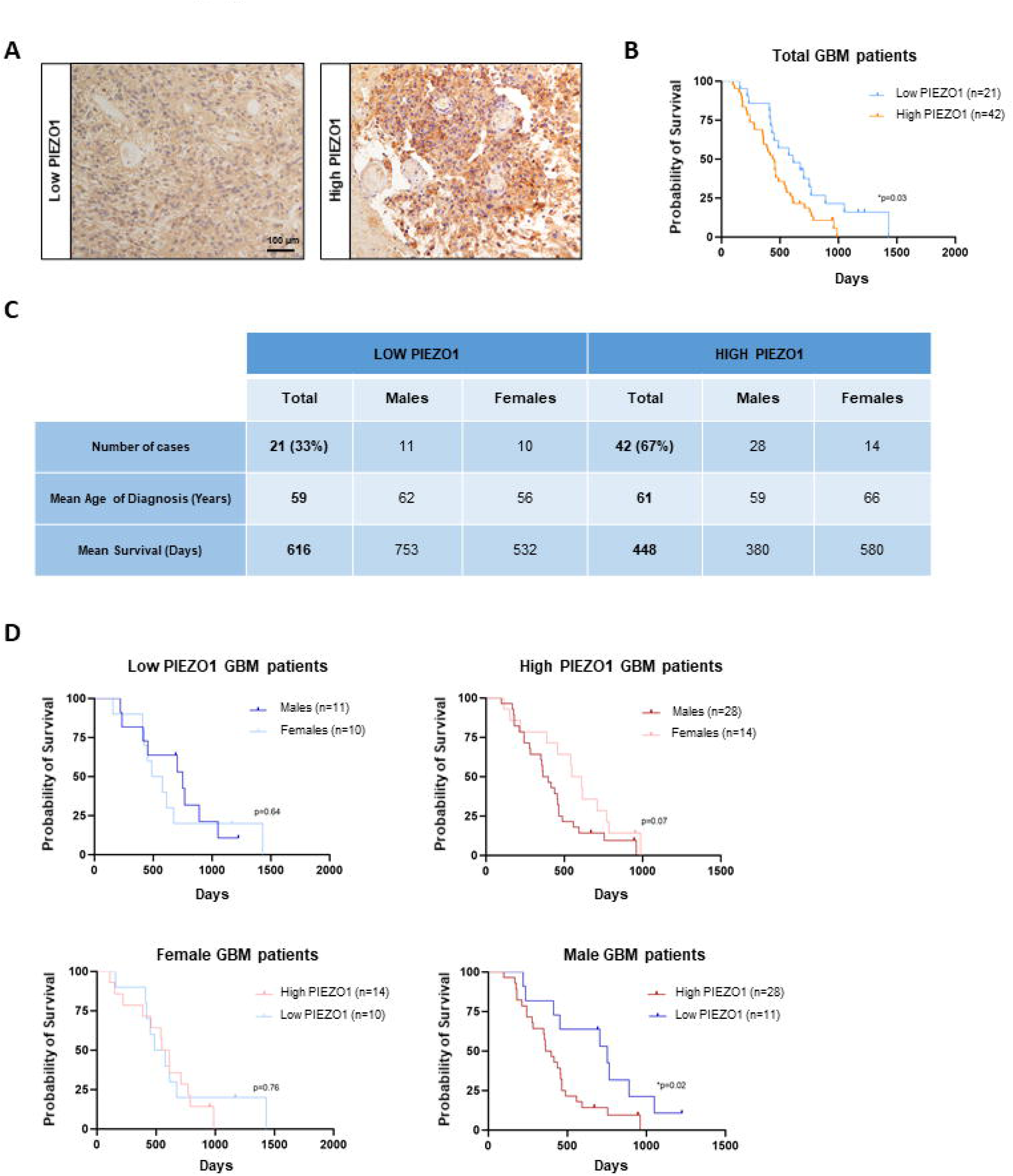
**(A)** Representative IHC images of PIEZO1-stained GBM human samples with high and low expression of this protein. **(B)** Kaplan-Meier survival curves of GBM patients classified according to their tumoral PIEZO1 expression levels. **(C)** Summary table of GBM patient data classified according to their tumoral PIEZO1 expression levels stratified by sex. **(D)** Kaplan-Meier survival curves of GBM patients classified according to their tumoral PIEZO1 expression levels and stratified by sex. To test differences between survival distributions, the log- rank (Mantel-Cox) test was used. Hazard ratios and confidence intervals were obtained by Mantel- Haenszel analysis.

Interestingly, PIEZO1 only behaved as a biomarker of worse prognosis in male patients, as no differences in survival were found in females. Likewise, while the male to female ratio (M:F) was 1:1 for low PIEZO1, it was 2:1 for high PIEZO1, indicating that there were more males than females with higher PIEZO1 levels (*Figure 1C-D*).

### Overexpression of PIEZO1 promotes a more aggressive phenotype in the U251 cell line

To explore the role of PIEZO1 in GBM biology, we genetically modified the U251 human GBM cell line with CRISPR-Cas9/SAM to generate an *in vitro* model of PIEZO1 overexpression (PIEZO1^OE^) (*Figure 2A-B*). As PIEZO1 is a Ca^2+^ channel, we first validated its functionality by studying the Ca^2+^ dynamics of these cells in a microfluidic system after the addition of the PIEZO1 activator Yoda1. As expected, we observed that intracellular Ca^2+^ levels peaked more rapidly in PIEZO1^OE^ cells than in control cells (transfected with the PIEZO1^EV^) (*Figure 2C*).

**Figure 2:**
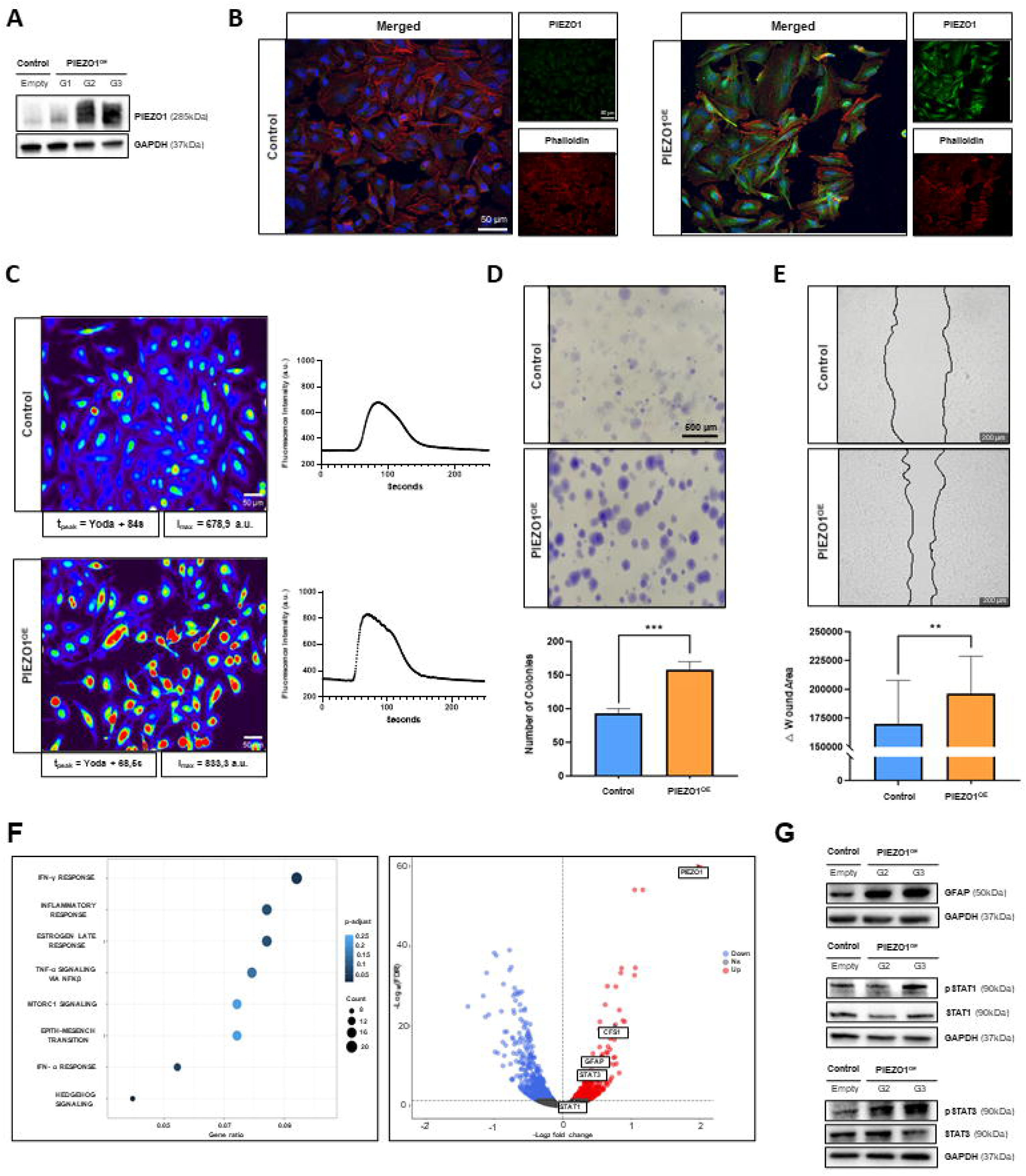
**(A)** Western blot showing PIEZO1 overexpression in U251 cells modified by CRISPR- Cas9/SAM employing three different sgRNAs (G1, G2, G3) compared to control (Empty) cells. **(B)** Representative immunofluorescence images showing PIEZO1 expression levels in control and PIEZO1^OE^ U251 cells. Phalloidin and DAPI were used as cytoskeleton and nucleus markers, respectively. **(C)** Representative live-cell images and quantification graphs of intracellular Ca^2+^ dynamics in control and PIEZO1^OE^ control cells treated with the PIEZO1 agonist Yoda1. Images showing the moment at which the intracellular Ca^2+^ peaked after Yoda1 addition. **(D-E)** Representative images and quantification graphs showing the differences in **(D)** clonogenicity and **(E)** migration between control and PIEZO1^OE^ U251 cells. **(F)** Graphical representation of the RNA-Seq results for control and PIEZO1^OE^ U251 cells. Left: Most significant pathways activated by PIEZO1 overexpression, as determined by GSEA. Right: Volcano plot highlighting activated inflammation-associated genes. **(G)** Western blot showing the activation of three astrocyte reactivity markers (STAT1 and STAT3 pathways, and GFAP) in PIEZO1^OE^ U251 cells. Quantitative variables are represented as the geometric mean ± SD. To compare two groups of quantitative variables, Student t-test was performed. **p<0.01; ***p<0.001.

After validating our *in vitro* model, we aimed to explore the role of PIEZO1 in GBM tumorigenesis. We found that PIEZO1 overexpression increased the clonogenicity (158±6.9 colonies in the PIEZO1^OE^ vs 93±11.8 in control, ***p<0.0001) and migration capacity (196224±32379 pixels of wound healing area in PIEZO1^OE^ vs 170093±37650 in control, **p=0.002) of U251 cells (*Figure 2D-E*), which are processes closely related to GBM aggressiveness and progression.

### PIEZO1 shifts U251 cells towards a reactive-like phenotype

To elucidate the molecular pathways underlying the phenotype observed in PIEZO1^OE^ U251 cells, we performed an RNA-Seq analysis of PIEZO1^OE^ and control cells. Gene Set Enrichment Analysis (GSEA) of the RNA-Seq results revealed that most of the pathways that were significantly enriched in PIEZO^OE^ cells compared to control cells were related to astrocyte reactivity and inflammation (*Figure 2F*). Thus, we validated these results by Western blotting. The GFAP, STAT1 and STAT3 molecules, which are three characteristic markers of astrocyte reactivity, were upregulated in PIEZO1^OE^ cells (*Figure 2G*).

### The overexpression of Piezo1 in astrocytes promotes glioma onset and reduces mouse survival *in vivo*

Given these *in vitro* findings, together with the fact that we observed higher PIEZO1 levels inside the tumour compared to the peritumoral area in the human GBM samples (*Figure 3A*), we wondered whether PIEZO1 could be a driver of glioblastoma *in vivo*. Thus, we generated a transgenic mouse colony that constitutively overexpressed murine Piezo1 under the astrocyte-specific promoter Gfap (*Piezo1*^Tg^/*Gfap*-*Cre*) (*Figure 3B*). We confirmed Piezo1 overexpression in the brains of these transgenic mice at different ages (*Figure 3C*).

**Figure 3:**
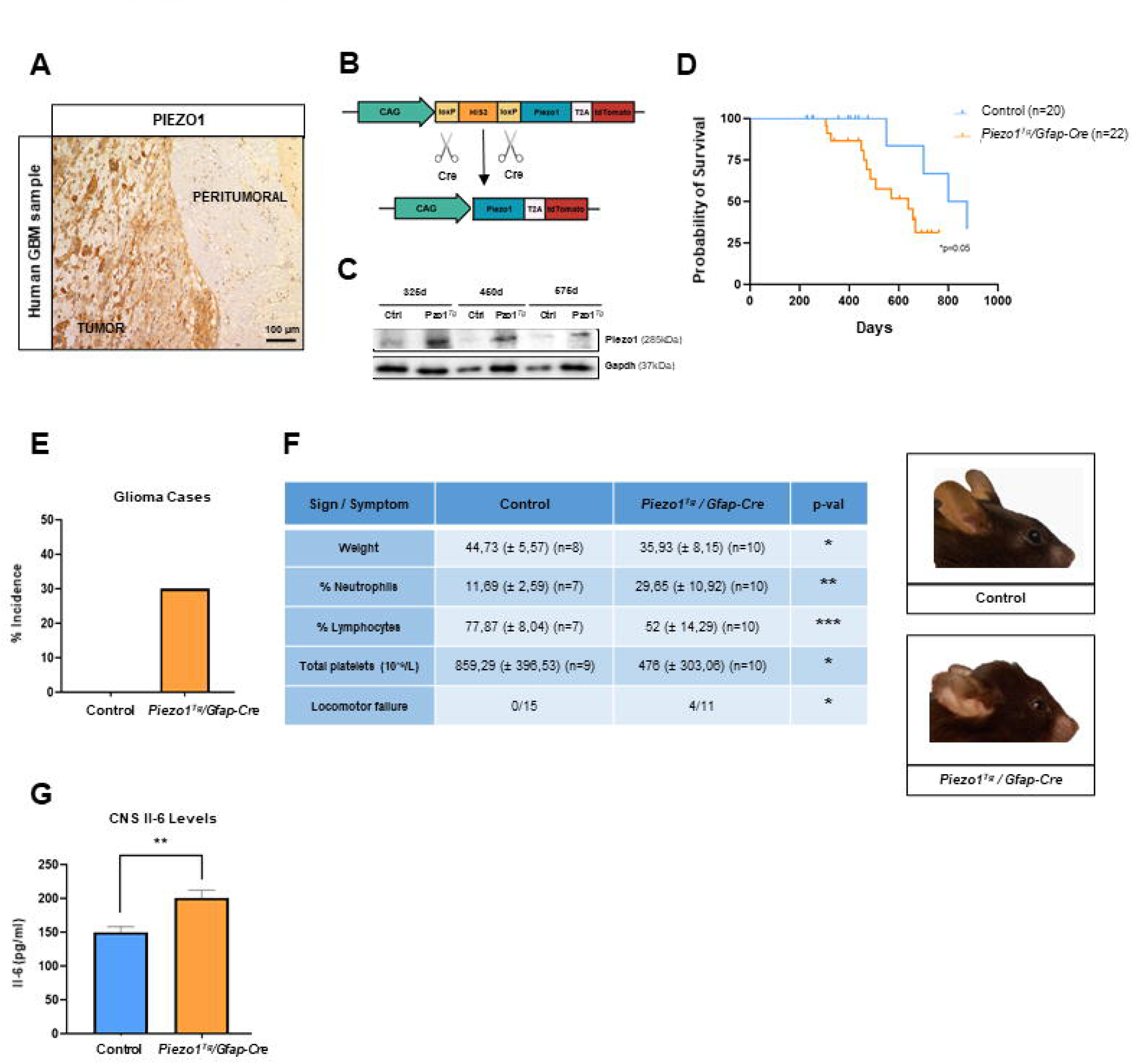
**(A)** Representative immunohistochemistry image of a PIEZO1-stained GBM human sample showing differences in the expression of this protein between the tumor and peritumoral areas. **(B)** Schematic representation of the function of the genetic system used to generate our *Piezo1*^Tg^/*Gfap-Cre* mouse model. **(C)** Western blot showing Piezo1 overexpression in *Piezo1*^Tg^/*Gfap-Cre* mouse brain samples when compared to age-matched control mouse brains. **(D)** Kaplan-Meier survival curves of *Piezo1*^Tg^/*Gfap-Cre* and control mice. **(E)** Graphical representation of the differences in the percentage incidence of glioblastoma between *Piezo1*^Tg^/*Gfap-Cre* and control mice. **(F)** Summary table and representative images of a series of signs and symptoms that significantly appeared in *Piezo1*^Tg^/*Gfap-Cre* mice that reached the humane endpoint compared to age-matched control mice. **(G)** Graphical representation of ELISA results showing a higher Il-6 concentration in nontumoral brain samples from *Piezo1*^Tg^/*Gfap-Cre* mice than in those from age-matched control mice. Quantitative variables are represented as the geometric mean ± SD. Qualitative variables are represented as n (cases) / n (total). To compare two groups of quantitative variables, Student t-test was performed. To compare two groups of qualitative variables, Fisher’s exact test was performed. To test differences between survival distributions, the log-rank (Mantel-Cox) test was used. Hazard ratios and confidence intervals were obtained via Mantel-Haenszel analysis. *p<0.05; **p<0,01; *p<0.001.

We observed that *Piezo1*^Tg^/*Gfap*-*Cre* mice presented a significant reduction in the overall survival compared to control mice (639 days in the *Piezo1*^Tg^/*Gfap*-*Cre* group vs 838 in the control group, *p=0.05) (*Figure 3D*). Interestingly, 30% of the transgenic mice developed glioma, a phenotype that was not observed in the control group (*Figure 3E*). *Piezo1*^Tg^/*Gfap*-*Cre* mice that reached the humane endpoint presented a series of significant phenotypic features such as weight loss, haematological alterations, locomotor failure and changes in head curvature (*Figure 3F*). To explore whether Piezo1 overexpression also promoted a reactive phenotype in astrocytes *in vivo*, similar to what was observed *in vitro*, we analysed Il-6 levels in non-tumoral mouse brain samples by ELISA and found that this proinflammatory cytokine was elevated in *Piezo1*^Tg^/*Gfap*-*Cre* brains (200.5±11.6 pg/ml in *Piezo1*^Tg^/*Gfap*-*Cre* brains vs 149.8±8.5 pg/ml in control brains; p=0.003) (*Figure 3G*).

### *Piezo1*^Tg^/*Gfap*-*Cre* mouse glioma present an aggressive phenotype

To characterize the phenotype of the gliomas developed in *Piezo1*^Tg^/*Gfap*-*Cre* mice, we performed histopathological analysis via immunohistochemical staining of a series of tumor markers. Interestingly, similar to human GBM samples, mouse gliomas presented higher expression of Piezo1 inside the tumor compared to the peritumoral zone (*Figure 4A*). As expected, tumor cells were positive for Ki67 and S100, proliferation and astrocytic markers, respectively (*Figure 4B*). We also observed positive staining for the glioblastoma-specific stemness markers Sox2 and Cd44 and for aggressiveness and reactive marker Stat3a (*Figure 4B*). As neuroinflammation is one of the most characteristic hallmarks of GBM, we evaluated the status of microglia and astrocytes in the peritumoral area. We observed high infiltration of both glial cell types in the tumor periphery, and these cells showed a clear reactive morphology (*Figure 4C*).

**Figure 4:**
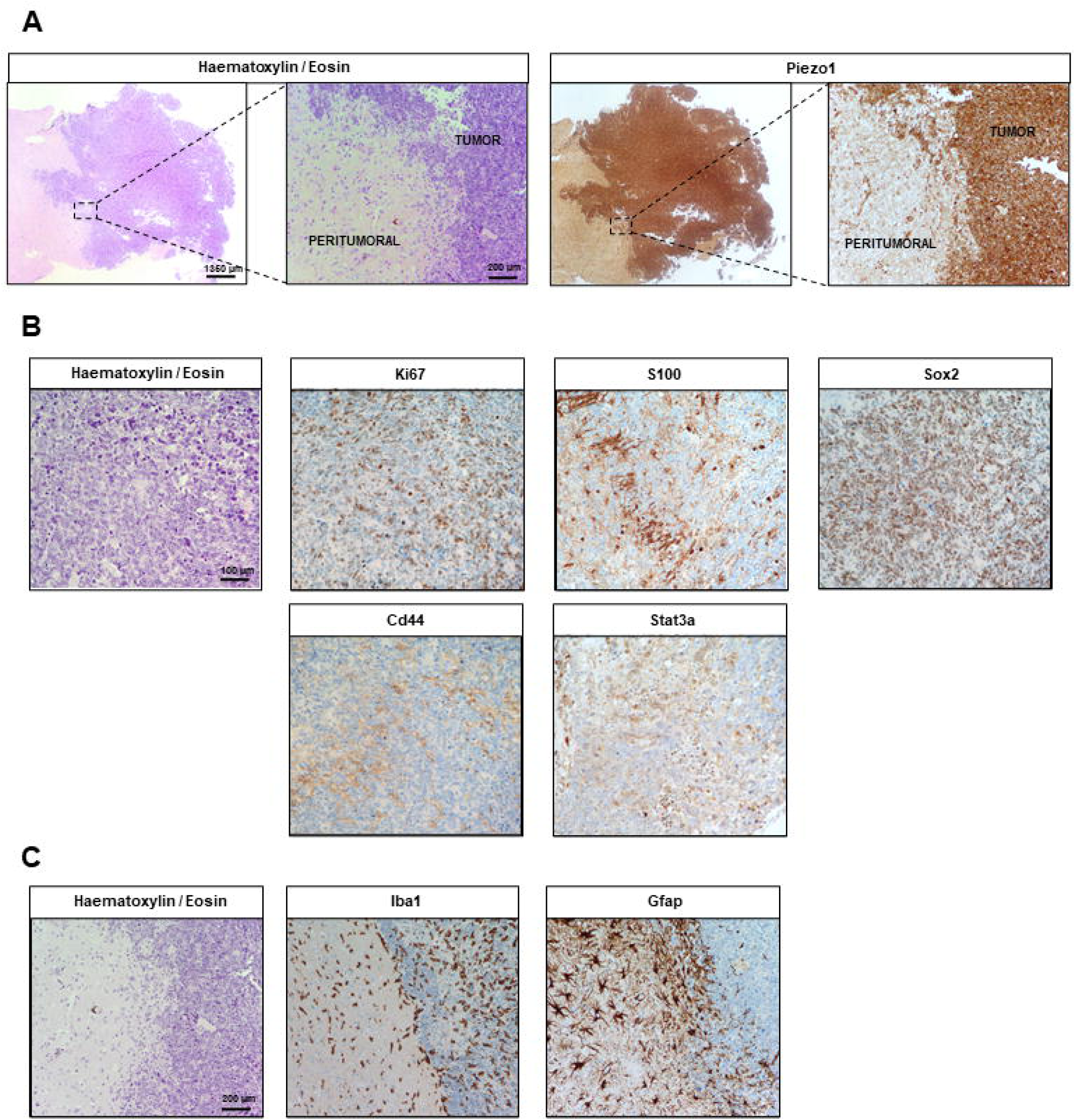
Representative immunohistochemistry images of a *Piezo1*^Tg^/*Gfap-Cre* mouse glioblastoma specimen. For each figure section, all images correspond to the same region and a reference haematoxylin/eosin staining image has been added. **(A)** Piezo1 staining showing differences in the expression of this protein between the tumoral and peritumoral areas. **(B)** Staining of different tumoral and glioblastoma-specific markers in an intratumoral region: Ki67 (proliferation); S100 (astrocytic); Sox2 and Cd44 (glioblastoma stemness); Stat3a (glioblastoma aggressiveness). **(C)** Staining of two glial markers in the tumor periphery region showing high reactive glial cell infiltration: Iba1 (microglia) and Gfap (astrocytes).

## Discussion

In this study, we demonstrated that PIEZO1 plays a key role in glioblastoma biology, both in humans and in our transgenic mouse model. We observed that Piezo1 overexpression in astrocytes leads to a significant reduction in overall survival elicited by glioblastoma development in 30% of transgenic mice. These observations are extremely relevant and could have an unprecedented impact on the glioblastoma paradigm since, to date, no gene with the capacity to generate gliomas on its own at this level of penetrance has been discovered. Therefore, Piezo1 could be the first described *bona fide* glioblastoma driver oncogene, and our study provides a new foundation for future glioma research. On the other hand, by generating a *de novo* transgenic mouse model of Piezo1 overexpression specifically in astrocytes (*Piezo1*^Tg^/Gfap-*Cre*), available for the scientific community, we offer a unique tool for researching the role of astrocytic Piezo1 in other CNS pathological processes, such as neurodegeneration^18^ and demyelination^19^. Furthermore, *Piezo1*^Tg^/*Gfap*-*Cre* is the first transgenic glioblastoma model on record that combines a single genetic modification and a remarkable tumour incidence, making it a highly effective research tool that faithfully reproduces the genomic and microenvironmental characteristics of human GBM, which could be used to greatly facilitate research on disease.

We hypothesize that the role of Piezo1 in gliomagenesis is related to the increase in intracellular Ca^2+^ levels caused by Piezo1 overexpression, as we observed in U251 PIEZO1^OE^ cells *in vitro*. Aberrant Ca^2+^ signals are a hallmark of reactive astrocytes, where there is an increase in Ca^2+^ wave amplitude, duration and/or frequency, although there is considerable variation in their spatiotemporal characteristics depending on the specific pathology^20–24^. Thus, high intracellular levels of Ca^2+^ in astrocytes are reported to promote reactivity, which is in agreement with the reactive-like phenotype observed in both our *in vitro* PIEZO1^OE^ U251 cell model and the *in vivo Piezo1*^Tg^/*Gfap*-*Cre* mouse model.

Consistent with these findings, we found that Il-6 levels in brain samples from *Piezo1*^Tg^/*Gfap*-*Cre* mice were significantly elevated compared to control mice. Neuroinflammation is one of the most relevant hallmarks of glioblastoma, and there is strong evidence that Il-6 plays a crucial role in both gliomagenesis and tumor progression through different mechanisms, especially through the activation of STAT3^25, 26^. STAT3 has been reported to be essential for proliferation and self-renewal in glioblastoma stem cells^27^, and its maintained activation promotes a chronic inflammatory state that favours cell malignant transformation^28^. Moreover, STAT3 activation in glioblastoma cells promotes the secretion of cytokines such as MIC-1 and TGF-β1, which favours the recruitment of immunosuppressive cells to the tumour microenvironment, thus eliciting immune evasion and promoting tumor growth^29, 30^. Interestingly, gliomas developed in *Piezo1*^Tg^/*Gfap*-*Cre* mice were positive for Stat3a marker, which is a prototypical marker of glioblastoma aggressiveness. These observations are in agreement with our previous findings that Piezo1 overexpression promotes reactivity and cytokine release in healthy astrocytes^18^. It is also well known that increased proliferation and migration are features of reactive astrocytes^31, 32^, which correlates with our *in vitro* results. Taken together, these results indicate that the chronic state of reactivity elicited by Piezo1 overexpression in astrocytes promotes phenotypic changes and maintains activation of inflammatory mediators, especially Stat3a, which are responsible for astrocyte malignancy.

Furthermore, when we analysed the status of glial cells surrounding gliomas of *Piezo1*^Tg^/*Gfap*-*Cre* mice, we observed that both microglia and astrocytes showed a clear reactive morphology, with elevated expression of the reactivity markers Iba1 and GFAP, respectively. Glioblastoma-associated astrocytes and microglia are reported to be immunosuppressive and to facilitate glioblastoma progression by promoting tumor immune evasion^33, 34^. This could indicate that Piezo1 overexpression in astrocytes not only promotes glioma onset but also favours tumour growth by generating an inflammatory response in the microenvironment. Nevertheless, the exact inflammatory mediators and intracellular pathways implicated in promoting gliomagenesis and tumor progression in Piezo1-overexpressing astrocytes need to be further investigated.

The data derived from our GBM patient cohort show that PIEZO1 also plays a key role in human glioblastoma. In agreement with previous reports^16, 17^, we observed that PIEZO1 is a marker of poor prognosis in GBM patients. Interestingly, we noted that PIEZO1 expression had an impact only on male patients, showing sex-related differences. PIEZO1 overexpression worsened GBM prognosis in males, while female overall survival did not differ between the low and the high groups. Furthermore, in the high PIEZO group, the M:F ratio was 2:1, and the mean survival was higher in women, which is in agreement with data reported for GBM patients worldwide^35^. Interestingly, in patients with low PIEZO1 expression, there were equal numbers of men and women, and the prognosis was better in males.

As we previously reported, the global incidence of GBM is nearly two times higher in males than females, and women have longer survival and better outcomes^36^. The sex-specific differences that lead to these observations have been deeply researched and discussed in the literature, and there are some cell-intrinsic sexual dimorphisms that could explain the higher impact of PIEZO1 overexpression on male GBM patients. Sun et al. reported that male astrocytes exhibit greater inactivation of the cell-cycle regulator retinoblastoma (RB) protein, which explains the predominance of GBM in men. It is well known that Ca^2+^ regulates RB protein inactivation by promoting its phosphorylation^37^. We hypothesize that this male-specific susceptibility to cell cycle changes is exacerbated by the maintained elevation of intracellular Ca^2+^ levels elicited by PIEZO1 overexpression, which could explain the sex-driven differences observed in our patient cohort. On the other hand, male astrocytes produce more proinflammatory cytokines associated with glioblastoma growth, such as IL-1β, IL-6, and TNF-α^38^. Given that our hypothesis is that PIEZO1 is directly implicated in astrocyte reactivity, its overexpression in male GBM cells could further increase these differences in cytokine secretion compared to that in female GBM cells, thus causing this worse prognosis unique to males in the high PIEZO1 group. Thus, although it would be necessary to increase the number of patients in our cohort and to further research the mechanisms implicated in the observed sex-driven differences, our data suggest that the classification of GBM patients according to their PIEZO1 tumoral expression levels could be useful in clinical practice to predict prognosis considering sex.

## Conclusion

Our study demonstrated that overexpression of the mechanoreceptor Piezo1 is sufficient to trigger the onset of glioblastoma, making it the first discovered *bona fide* glioblastoma driver oncogene. We also observed that high tumoral expression of PIEZO1 is associated with a worse prognosis in male GBM patients, suggesting that PIEZO1 is a new biomarker for this disease. Furthermore, we developed a novel and unique transgenic mouse model of Piezo1 overexpression specifically in astrocytes (*Piezo1*^Tg^/*Gfap*-*Cre*), which could be used by the scientific community to study the role of Piezo1 not only in glioblastoma but also in other neurological processes and diseases.

Although the mechanism by which Piezo1 promotes gliomagenesis needs to be further investigated, we hypothesize that the overexpression of this mechanoreceptor promotes Ca^2+^ flux and signalling that causes chronic reactivity of astrocytes, which triggers their malignancy and generates a neuroinflammatory microenvironment that favours tumor progression, thus identifying Piezo1 as a potential therapeutic target for GBM.

## Acknowledgements

We would like to thank the Transgenic Mouse Editing Unit at CNIO for their assistance at generating the transgenic model; the Animal Facility Unit staff at CNIO for their assistance in the breeding of the mouse colony; the Histopathology Unit at CNIO for their assistance at mouse IHC; and Dr. Massimo Squatrito for his generous transfer of the U251 cell line. This project was funded by the European Union’s Horizon 2020 Research and Innovation Program under the Marie Sklodowska-Curie grant agreement No 101027864 awarded to M.V.E.; by Instituto de Salud Carlos III through the funding "CP19/00140" and project "PI18/00295" (cofunded by the European Regional Development Fund/European Social Fund "A way to make Europe"/"Investing in your future") awarded to M.G.; by Fundación Tatiana Pérez Guzmán del Bueno through a PhD fellowship awarded to P.B.C; by Instituto de Salud Carlos III through the funding "CD19/00222 (Cofunded by European Regional Development Fund/European Social Fund "A way to make Europe"/"Investing in your future" awarded to M.H.S.; by Instituto de Salud Carlos III (ISCIII) through the grant "FI22/00234" and cofunded by the European Union awarded to A.O.S.; and by the CRIS contra el Cancer Foundation, CRIS-CNIO agreement 2017-2020, and CRIS-CNIO agreement 2020-2023.

## Author contributions

M.V.E. conceived and planned the study with input from M.G and J.M.L. M.V.E, M.G, P.B.C, MA.N.A, designed and performed the cultures, cellular and molecular biology experiments and *in vivo* experiments. M.H.S designed the guides for CRISPR-Cas9 experiments. S.O. designed the transgenic murine model. R.G, RM.G.M, A.H.L obtained patient samples and clinical information, and performed the IHC experiments on human samples. O.D. performed the RNA-seq while A.A.B and O.G analysed the data. M.P.M designed the microfluidic experiment and M.P.M and J.G.A performed the confocal experiments with M.V.E and P.B.C. M.V.E, M.G. and J.M.L supervised the study. P.B.C wrote the manuscript with contributions from M.V.E and M.G. M.V.E. edited the final manuscript. All authors approved the final manuscript.

## Methods

### Human brain samples

Human brain biopsy samples from glioblastoma patients were obtained from the Hospital 12 de Octubre de Madrid (Madrid, Spain), in accordance with the Declaration of Helsinki and the approval of the Ethics Committee of the Hospital 12 de Octubre (CEIm number 21/161). Written informed consent was obtained from all patients.

A cohort of 63 patients was collected. Immunohistochemistry was performed on 3µm thickness sections of formalin-fixed and paraffin-embedded brain samples. Immunostaining was performed on a Leica Bond-III stainer platform (Leica Biosystems, UK) using the Leica Bond Polymer Refining kit (Leica Biosystems), heat mediated antigen retrieval with Tris-EDTA buffer (pH 9.0) for 30 min following incubation with 1:100 dilution of the primary antibody anti- Piezo1 (28511- 1-AP, Proteintech). Samples were scored in five levels of PIEZO1 expression in the tumor (0 to 4), using the PIEZO1 staining intensity in the healthy area of each sample as a reference. Groups 0 and 1 correspond to a tumor staining intensity lower and similar to that of the healthy area, respectively, and were classified as “low PIEZO1 expression”. Levels 2 to 4 correspond to tumor PIEZO1 expression higher than in the non-tumoral area, and were classified as “high PIEZO1 expression”.

### Generation of transgenic Tg.CAG-LSL-Piezo-T2A-Tomato mice

The transgenic construct was assembled in the plasmid p15A by the company Gen-H Genetic Engineering Heidelberg GmbH. For mouse production, DNA was prepared from a bacterial extract using the EndoFree Plasmid Maxi Kit (#12362, Qiagen) and digested with MluI to release the plasmid backbone. The band containing the construct (13,200 bp) was gel-purified (0.8% agarose gel) using the QIAquick Gel Extraction Kit (#28704, Qiagen) and diluted in microinjection buffer containing 10 mM Tris-HCl (pH 7.8) and 0.1 mM EDTA at 1 ng/µl. Transgenic mice were generated by pronuclear injection of the transgenic DNA into B6.CBA x B6 zygotes obtained from superovulated B6.CBA females crossed with C57Bl6 males using standard protocols^39^. Among the 28 mice born, 3 were confirmed to be positive for complete transgene integration. Founders 1 and 2 were crossed with C57Bl6 females to test for germ line transmission. This line, *Piezo1*^Tg^, was maintained by crossing hemizygous animals (T/+) with wild-type C57Bl6 mice.

To generate the mouse colony of study, *Piezo1*^Tg^ mice were crossed with a mouse strain carrying astrocyte-specific expression of Cre recombinase, *Gfap-Cre* mice, to generate *Piezo1*^Tg^/*Gfap-Cre* mice. All animals were maintained at the CNIO Animal Facility under specific pathogen-free conditions in accordance with the recommendations of the Federation of European Laboratory Animal Science Associations (FELASA). The animals were treated following the Principles of Laboratory Animal Care (EU 609/86 CEE, Spanish Real Decreto 223/88 BOE-18/03) and guidelines for the care of experimental animals of the CNIO. The generation of the mouse model and all animal experimentation were approved by the Institutional Animal Care and Use Committee (IACUC) and the Ethical Committee (CEIyBA), under the PROEX 265.0/21.

### Mouse histopathological analysis and immunohistochemistry

Brain samples were fixed in 10% neutral buffered formalin (4% PFA in solution), paraffin-embedded, cut at 3µm thickness, mounted on TOMO slides and dried overnight. Before staining, the slides were deparaffinized in xylene and rehydrated through a series of graded ethanol until water. Consecutive sections were stained with haematoxylin and eosin (H&E), and several immunohistochemistry reactions were performed on automated immunostaining platforms (Autostainer Link 48, Dako; Ventana Discovery XT, Roche).

Antigen retrieval was first performed with the appropriate pH buffer (low or high pH buffer, Dako; CC1m Ventana, Roche), and endogenous peroxidase was blocked (peroxide hydrogen at 3%). Then, the slides were incubated with the appropriate primary and secondary antibodies as detailed in *Table 1*. Immunohistochemical reactions were developed using 3,30-diaminobenzidine tetrahydrochloride (DAB) (Chromo Map DAB, #760-159, Ventana, Roche) and nuclei were counterstained with Carazzi’s haematoxylin. Finally, the slides were dehydrated, cleared and mounted with permanent mounting media for microscopic evaluation. Positive control sections known for the primary antibody were included for each staining run.

**Table 1:**
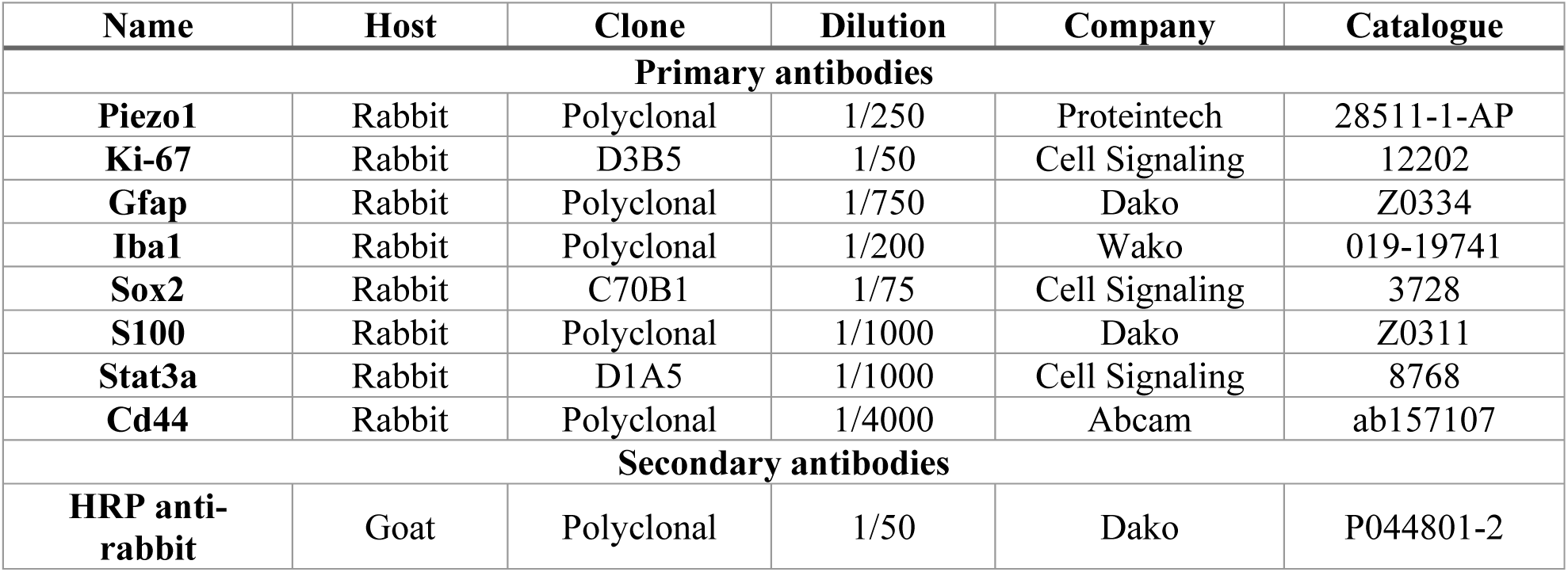
IHC antibodies.

### Enzyme-linked immunosorbent assay (ELISA)

Non-tumoral brain tissue samples from euthanized mice were collected and lysed in RIPA lysis buffer (#20-188, Millipore) containing protease inhibitors (Complete Mini, #11836153001, Roche) and phosphatase inhibitors (PhosSTOP, #4906837001, Roche) by sonication. Levels of Il-6 were determined using a mouse Il-6 DuoSet ELISA Kit (DY406, R&D) following the manufacturer’s instructions.

### Generation of PIEZO1-overexpressing human glioblastoma cells

To generate PIEZO1-overexpressing (PIEZO1^OE^) human glioblastoma cells, we transduced U251 cells (#09063001, Sigma Aldrich) with lentiviral particles containing the plasmids lentiMPHv2 (#89308, Addgene) and lentiSAMv2 (#75112, Addgene) containing the sgRNAs to activate *PIEZO1*. These guides were designed using the online CRISPR design tool (http://crispr-era.stanford.edu/) and were cloned and inserted into the plasmid lentiSAMv2 as previously described^40, 41^. The correct insertion of the sgRNA sequences was confirmed by Sanger sequencing. The empty plasmid lentiSAMv2 without a targeting sgRNA was used as a control. Lentiviral particle production was performed as previously described^42^. Lentivirus were used to infect U251 cells in medium supplemented with 8 µg/ml polybrene. Cells were selected with 2 µg/ml blasticidin (#B-1247, AG Scientific) and 400 µg/ml hygromycin (#10843555001, Roche) for 7 days. The sequences of the sgRNAs used were as follows: PIEZO1_sgRNAact_TSS11 (guide 1): 5’- CACCGTTATAAAGGCCCGCGGGCGG – 3’ PIEZO1_sgRNAact_TSS76 (guide 2): 5’ – CACCGCCGCCTCCGCGCTTCCCCGA – 3’ PIEZO1_sgRNAact_TSS143 (guide 3): 5’ – CACCGAGGCCCCAACGCACCAGGGC – 3’ Modified U251 cells were cultured in Dulbecco’s modified Eagle’s medium (DMEM; D5796, Sigma), supplemented with 10% foetal bovine serum (FBS; F7524, Sigma), 5 µg/ml penicillin/streptomycin (P/S; SOPENSTREP100ML, Solmeglas), and 15mM HEPES (83264, Sigma), and maintained at 37°C and 5% CO_2_. These cells are referred to as *PIEZO1^SAM^* U251 cells (*PIEZO1^EV^* if empty vector, *PIEZO1^OE^*if overexpression)

### RNA-Seq analysis of Piezo1-overexpressing U251 cells

Total RNA samples from *PIEZO1^SAM^* U251 cells [1µg; RQS/RIN = 9.3, range 9.0-9.5] were converted into sequencing libraries with the “NEBNext Ultra II Directional RNA Library Prep Kit for Illumina” (#E7760, New England Biolabs) following the manufacturer’s instructions. Briefly, polyA+ fraction is purified and randomly fragmented, converted to double stranded cDNA and processed through subsequent enzymatic treatments of end-repair, dA-tailing, and ligation to adapters. Adapter-ligated library is completed by PCR with Illumina PE primers. This kit generates directional libraries stranded in the antisense orientation [the read1 (the only read in single read format) has the antisense orientation]. The resulting purified cDNA libraries were applied to an Illumina flow cell for cluster generation and sequenced on an Illumina NextSeq 500 (with v2.5 reagent kits) following manufacturer’s protocols.

The resulting single-end reads (86 bases) were analysed with the nextpresso pipeline^43^ as follows: Sequencing quality was checked with FastQC v0.11.0 (https://www.bioinformatics.babraham.ac.uk/projects/fastqc/). Reads were aligned to the human genome (GRCh38) with TopHat2^44^ using Bowtie1^45^ and SAMtools^46^, allowing 3 mismatches and 20 multihits. The Gencode v41 gene annotation for GRCh38 was used. Read counts were obtained with HTSeq^47^. Differential expression and normalization were performed with DESeq2^48^ keeping only those genes where the normalized count value was higher than 10 in at least 30% of the samples. Finally, those genes that had an adjusted p-value below 0.05 FDRs were selected. GSEAPreranked was used to perform gene set enrichment analysis for the selected gene signatures on a preranked gene list, setting 1000 gene set permutations^49^. Only those gene sets with significant enrichment levels (FDR q-value < 0.25) were considered.

The data discussed in this publication have been deposited in the NCBI Gene Expression Omnibus^50^ and are accessible through GEO Series accession number GSE242946 (https://www.ncbi.nlm.nih.gov/geo/query/acc.cgi?acc=GSE242946).

### Live-cell calcium (Ca^2+^) imaging

Cells were seeded onto µ-Slide VI 0.4 (#80606, Ibidi) channels and grown for 24h until 70% confluence. The cells were then incubated with 3 µM Cal-520AM dye (ab171868, Abcam) in Hank’s balanced salt solution (HBSS; no phenol red) supplemented with 10mM glucose and 25mM HEPES for 90 min at 37°C, followed by 30 min at room temperature. Cells were protected from bright light at all times.

Time-lapse Ca^2+^ imaging was performed using a DMI6000B fluorescence microscope from Leica (Leica Microsystems) equipped with a 20X NA0.5 dry objective, L5 fluorescence filtercube, a Hamamatsu-ORCA-ER camera and LAS AF v2.7 software. Videos were capture with a 4fps rate and a 4x4 binning. The flow rate was 0.39ml/min corresponding to a physiological blood vessel shear stress of 0.5dyn/cm^2^ or 0.79ml/min for a shear stress of 1dyn/cm^2^. The syringe pump used was a NE-1600 syringe pump model (New Era Pump Systems Inc). Following a 30s baseline, cells were stimulated with 5µM Yoda1 for 150s.

### Immunofluorescence

Cells were fixed with 4% PFA in PBS for 10 minutes at room temperature (RT), permeabilized and blocked at the same time with blocking buffer (0.5% BSA, 0.1% Triton X-100 in 1X PBS) overnight at 4°C. After this, the samples were incubated with 1:150 dilution of Piezo1 primary antibody (1:150, Proteintech, 28511-1-AP) in blocking buffer, overnight at 4°C. After washing with PBS, the following secondary antibody was added overnight at 4°C in blocking buffer: Alexa Fluor^TM^ 647 donkey anti-rabbit IgG (1:1000, Invitrogen #A-32795). Finally, cell nuclei and cytoskeleton were stained with 0.5 µg/mL DAPI (Invitrogen D1306) and Phalloidin Alexa Fluor^TM^ 568 (Invitrogen A12380) diluted in water for 15 minutes at room temperature. Images were acquired using a Leica SP5 confocal microscope.

### Migration assay

Cells were seeded onto 12-well plates (#30012, SPL Life Sciences) and grown to 90% confluence. “Wound” gaps were generated as a longitudinal scratch of the well using a 20 µl pipette tip. Six different areas of the wound per well were imaged using a Leica Thunder microscope. Images were automatically taken every 10 min for 24h, using a 10x magnification objective. The width of the area containing no cells (wound area) was calculated using ImageJ software (http://imagej.nih.gov/ij), and the Δ wound area was calculated as the difference between the initial and final wound areas.

### Colony formation assay

Cells were seeded onto 6-well plates (#30006, SPL Life Sciences) at a density of 500 cells/well and grown for two weeks on complete media. Then, the colonies were fixed with 100% methanol for 10 min and stained with 0.1% crystal violet (#C0775, Sigma Aldrich) for 30 min. After three washes with PBS, the plates were allowed to air dry, and the number of colonies was counted.

### Western-blot

Western blotting was performed using protein lysates from *PIEZO1*^SAM^ U251 cells and *Piezo1*^Tg^ mouse brains. Tissues and cells were homogenized in RIPA lysis buffer (#20-188, Millipore) containing protease and phosphatase inhibitors (#11836153001, Complete Mini, Roche; and #4906837001 PhosSTOP, Roche, respectively). Soluble proteins were boiled in Laemmli buffer (#1610747, Bio-Rad) supplemented with β-mercaptoethanol, resolved on a 6-15% gradient SDS‒ PAGE gel, and transferred to a PVDF membrane. The membranes were blocked in 5% non-fat milk for 1 hour at room temperature and then incubated with primary antibodies in bovine serum albumin (BSA; A7906, Sigma) Tween-Tris buffered saline (T-TBS) solution, overnight at 4°C.

After primary antibody incubation, membranes were washed in T-TBS and incubated with the corresponding HRP-conjugated secondary antibody for detection by the enhanced chemiluminescence SuperSignal® West Femto Maximum Sensitivity Substrate (#34095, Thermo Scientific). GAPDH was used as a cellular loading control. Antibodies used are listed in *Table 2*.

**Table 2:** WB antibodies.

| Name | Host | Clone | Dilution | Company | Catalogue |
| --- | --- | --- | --- | --- | --- |
| **Primary antibodies** |  |  |  |  |  |
| **PIEZO1** | Rabbit | Polyclonal | 1/300 | Proteintech | 28511-1-AP |
| **GFAP** | Mouse | GA5 | 1/1000 | Cell Signaling | 3670 |
| **STAT1** | Rabbit | 42H3 | 1/1000 | Cell Signaling | 9175 |
| **pSTAT1 (Tyr701)** | Rabbit | 58D6 | 1/1000 | Cell Signaling | 9167 |
| **STAT3** | Mouse | 124H6 | 1/1000 | Cell Signaling | 9139 |
| **pSTAT3 (Tyr705)** | Rabbit | D3A7 | 1/1000 | Cell Signaling | 9145 |
| **GAPDH** | Rabbit | 14C10 | 1/1000 | Cell Signaling | 2118 |
| **Secondary antibodies** |  |  |  |  |  |
| **HRP anti-rabbit** | Goat | Polyclonal | 1/5000 | Dako | P0448 |
| **HRP anti-mouse** | Goat | Polyclonal | 1/5000 | Dako | P0447 |

### Statistical analysis

All the statistical analyses were performed using GraphPad Prism 10 (GraphPad Prism^®^). The normality of the data was assessed using the D’Agostino criteria before any further statistical tests were performed. When two groups of quantitative variables were compared, Student’s t-test or Mann-Whitney test was performed. To compare two groups of qualitative variables, Fisher’s exact test was performed. To test differences between survival distributions, the log-rank (Mantel-Cox) test was used. Hazard ratios (HRs) and confidence intervals (CIs) were obtained via Mantel-Haenszel analysis. A *p* value ≤ 0.05 was considered to indicate statistical significance. Further details of the statistical analysis performed are given in each figure legend.

---

**Supplemental Figure 1S:**
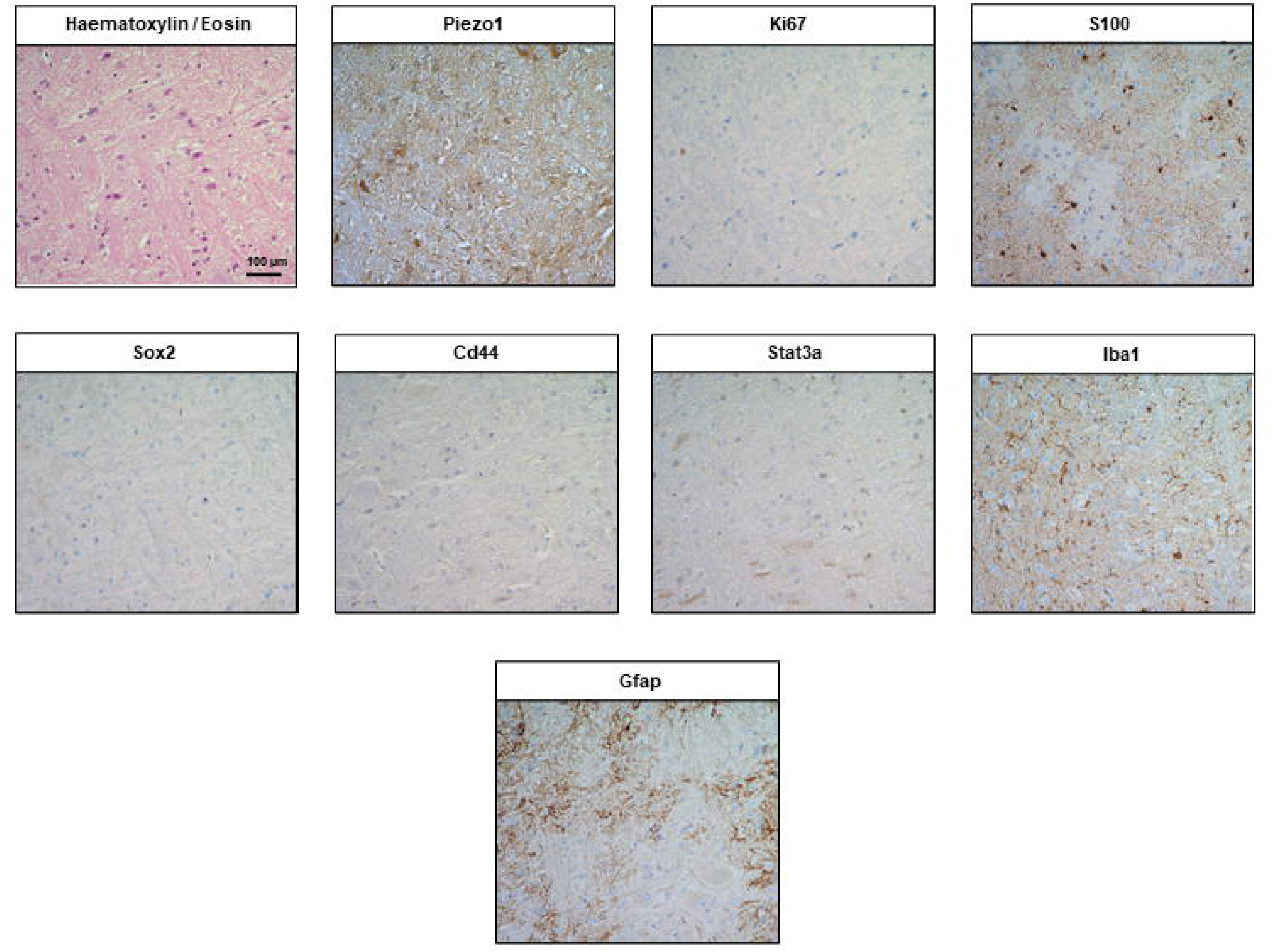
Representative immunohistochemistry images from a healthy brain from a control mouse age-matched with a *Piezo1*^Tg^/*Gfap-Cre* glioblastoma mouse from Figure 4. All images correspond to the same region and a haematoxylin/eosin staining image has been added as a reference. Staining markers are the same shown for the glioma sample in Figure 4.

